# The contribution of area MT to visual motion perception depends on training

**DOI:** 10.1101/094128

**Authors:** Liu D. Liu, Christopher C. Pack

## Abstract

**Summary:** Perceptual decisions require the transformation of raw sensory inputs into cortical representations suitable for stimulus discrimination. One of the best-known examples of this transformation involves the middle temporal area (MT) of the primate visual cortex. Area MT provides a robust representation of stimulus motion, and previous work has shown that it contributes causally to performance on motion discrimination tasks. Here we report that the strength of this contribution can be highly plastic: Depending on the recent training history, pharmacological inactivation of MT can severely impair motion discrimination, or it can have little detectable influence. Similarly, depending on training, microstimulation can bias motion perception or simply introduce noise. Further analysis of neural and behavioral data suggests that training shifts the readout of motion information between MT and lower-level cortical areas. These results show that the contribution of individual brain regions to conscious perception can shift flexibly depending on sensory experience.

## Introduction

Standard models of perceptual decision-making revolve around a specific set of operations. The first concerns the transformation of sensory signals into a representation that facilitates task performance. The second involves integrating the output of these transformations over space and time, after which the corresponding motor response can be initiated (Gold and Shadlen, 2007; Mazurek et al., 2003). Probably the best-known example of these operations is in the domain of visual motion perception: Estimates of motion are computed by populations of neurons in visual cortex, with the integration of information occurring in downstream areas of the parietal and frontal cortices (Cook and Maunsell, 2002; Gold and Shadlen, 2000; Katz et al., 2016; Kim and Shadlen, 1999; Platt and Glimcher, 1999; Roitman and Shadlen, 2002; Shadlen and Newsome, 2001).

In this context, the middle temporal visual area (MT) is thought to play a special role by transforming low-level representations of motion into a representation that is both sharply direction-selective and invariant to other stimulus features (Born and Bradley, 2005; Simoncelli and Heeger, 1998). Consistent with this idea, MT lesions severely impair performance on motion discrimination tasks (Hess et al., 1989; Newsome and Pare, 1988; Rudolph and Pasternak, 1999), and electrical microstimulation of MT can strongly bias the animals’ choice of motion direction (Salzman et al., 1990; Salzman et al., 1992). Thus MT activity appears to be both necessary and sufficient for accurate discrimination of motion direction.

These conclusions have emerged from previous studies in which the motion stimulus was corrupted by noise, the most common example being a moving fields of random dots. In this stimulus, a subset of the dots moves in a coherent direction, with the rest moving in random directions (Britten et al., 1992; Newsome and Pare, 1988); the subject’s task is to recover the direction of the coherent dots. MT is similarly critical for the discrimination of moving grating stimuli corrupted with noise (Rudolph and Pasternak, 1999). At the single-cell level, individual MT neurons exhibit clear selectivity for motion even in the presence of large amounts of noise (Britten et al., 1992), but they are also selective for other stimuli such as gratings (Movshon et al., 1985), bars (Pack and Born, 2001), and naturalistic movies (Nishimoto and Gallant, 2011). The formation of such invariant stimulus representations is considered on theoretical grounds to be an essential step in perceptual decision making (DiCarlo and Cox, 2007; Tsunada et al., 2016). On the other hand, there are many other visual cortical areas with neurons tuned to motion direction (Cook and Maunsell, 2002; Desimone and Ungerleider, 1986; Fattori et al., 2009; Felleman and Van Essen, 1987; Gegenfurtner et al., 1997; Law and Gold, 2008; Levitt et al., 1994; Movshon et al., 1985; Peterhans and von der Heydt, 1993), and in principle any one of these could support perceptual decision-making, for appropriate choices of stimuli.

In this regard, drifting grating stimuli provide an interesting test, as they elicit strong direction selectivity in subpopulations of neurons in lower-level cortical areas such as V1, V2, and V3, as well as MT (El-Shamayleh and Movshon, 2011; Felleman and Van Essen, 1987; Levitt et al., 1994; Liu et al., 2016; Movshon et al., 1985). Although monkeys are capable of discriminating such stimuli following recovery from MT lesions (Pasternak and Merigan, 1994; Rudolph and Pasternak, 1999), little is known about the neural basis of such discriminations in intact animals. We therefore trained monkeys to discriminate the motion of these stimuli, and perturbed MT activity using reversible inactivation and microstimulation. These experiments, along with data from single-neuron recordings, yielded the surprising result that the role of MT in motion perception depends strongly on recent perceptual experience: After several sessions of training with grating stimuli, we found that MT played little or no discernable role in motion perception. However, after training with random-dot stimuli, MT became strongly and causally associated with the perception of the same grating stimuli. These results support the idea that the causal connection between cortical activity and conscious perception is highly plastic (Chen et al., 2016; Chowdhury and DeAngelis, 2008).

## Results

We studied the relationship between MT activity and perception in the context of two different training periods. In the first period, monkeys spent several weeks performing a discrimination task in which the stimulus was a drifting grating (**Figure 1A**). This stimulus elicits strongly direction-selective responses in many visual cortical areas, including V1, V2, V3, and MT (Felleman and Van Essen, 1987; Gegenfurtner et al., 1997; Levitt et al., 1994; Movshon et al., 1985; Peterhans and von der Heydt, 1993). In the second period, the same animals were trained on a task requiring motion discrimination for random-dot patches. The representation of motion for such stimuli appears to rely heavily on MT (Hess et al., 1989; Newsome and Pare, 1988), and it is far less reliable in V1 (Hammond and MacKay, 1977; Skottun et al., 1988). Following each training period, we examined the link between MT activity and motion perception using single-unit recordings, pharmacological inactivation, and electrical microstimulation.

**Figure 1.**
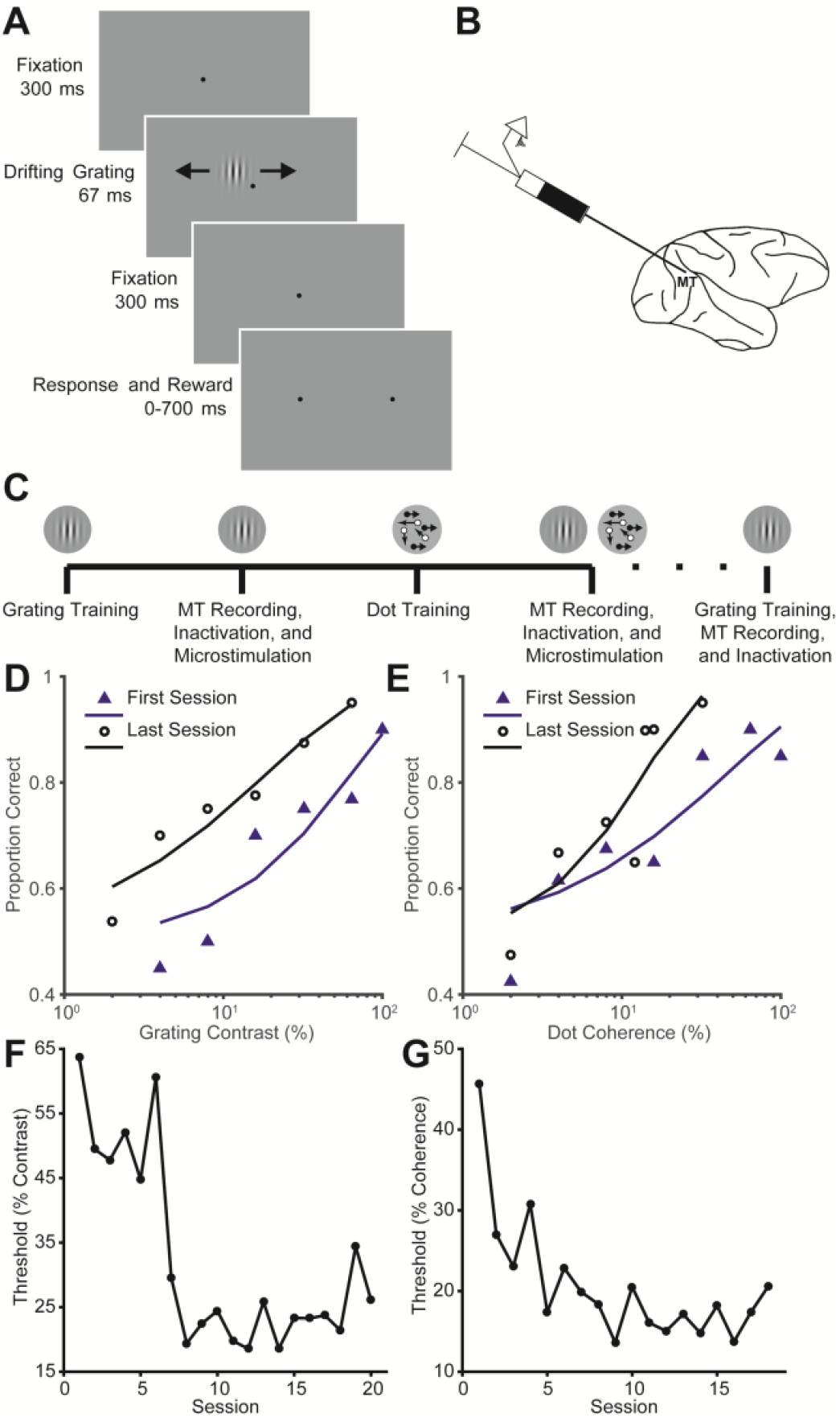
Discrimination tasks and sequence of training. **(A)** The animals were required to maintain fixation in a 2° window for 300 ms before the onset of the stimulus. The Gabor patch was centered in the envelope of the receptive fields of the units recorded. After the offset of the stimulus, the animals were required to fixate for another 300 ms until the fixation spot disappeared and to initiate a response within 700 ms. A reward was dispensed if the animal made an eye movement to the peripheral target that corresponded to the direction of stimulus motion. **(B)** Schematic illustration of the experimental setup. We recorded neural activity using a multi-channel probe, with a built-in cannula for muscimol injections. The microstimulation experiments used single contact platinum-iridium electrodes for recording and stimulation. **(C)** We trained naïve monkeys first on grating motion discrimination, followed by recording, reversible inactivation, and microstimulation of MT to test its causal contribution to motion perception. We then trained the monkeys on random dots motion discrimination, followed by the same recording, inactivation, and microstimulation protocols to test MT’s contribution to both grating and dots motion. In one monkey, we again trained the animal repeatedly on the grating discrimination task, with no further exposure to the random dots. MT’s contribution to grating motion was tested with inactivation. **(D)** Example psychometric functions for grating motion discrimination before and after training. The first session is defined as the first session of performance after the animal achieved as least 75% correct on the 100% contrast condition. The last session is the last session of performance before the experiments began. 40 trials were performed at each stimulus condition. Lines represent fits to Weibull functions using the maximal likelihood procedure. **(E)** Example psychometric functions for random dot stimuli before and after training. **(F)** The psychophysical contrast threshold as a function of the number of sessions for the grating motion discrimination. Threshold was defined as 92% correct of the Weibull function fits. **(G)** The psychophysical coherence threshold as a function of the number of sessions for the random-dot motion discrimination task.

## Contribution of MT to motion perception after training on grating stimuli

Both animals were initially trained to indicate the direction of motion for a briefly presented, drifting Gabor grating (**Figure 1A**). We manipulated task difficulty by varying the contrast of the Gabor patch, and over the course of several weeks, the threshold for accurate (92%) performance plateaued at around 20% contrast (**Figure 1F**), as found previously for brief stimuli (Tadin et al., 2003). We then recorded from MT, using linear electrode arrays.

As expected from much previous work (Albright, 1984; Khawaja et al., 2009; Movshon et al., 1985), individual MT neurons were well tuned for grating motion, and there was a small but significant correlation between firing rate fluctuations and behavioral choices (Britten et al., 1996). This correlation can be quantified with choice probability (CP), and we found that CP values were on average slightly greater than chance (0.515 ± 0.059, permutation test; *P* = 0.03).

Despite this correlation, we found that reversible inactivation of MT had little effect on behavioral performance after training with grating stimuli (**Figure 2C** **and** **F**). Specifically, we injected muscimol into the area of MT that represented the location of the stimulus (see Methods) and verified that neuronal activity in the vicinity of the electrode was silenced (**Figure S1**). Previous work has shown that injecting the same volume of muscimol into MT leads to profound deficits in motion perception that peak at 18 hours after the injection (Chowdhury and DeAngelis, 2008); at this time point the muscimol has typically spread over a radius of ∼1-2 mm. However, we found little evidence of a behavioral deficit in either monkey at any time period (**Figure 2C** **and** **F**). When animals were tested at 18 hours after the injection, behavioral thresholds had increased by an average of only 0.6 ± 5.0% (2.1 ± 3.6% for monkey Y and -2.8 ± 6.7% for monkey C). This change was comparable to that of the variability found in the absence of inactivation, and it was not significantly different from the pre-injection baseline (Wilcoxon rank sum test; *P* = 0.24; *P* = 0.07 for monkey Y and *P* = 0.84 for monkey C). Similar results were obtained with psychophysical testing performed at 45 minutes and at 48 hours after the injection (**Figure S6C, F**). Thus while MT activity was correlated with perceptual decisions about grating motion, we did not find evidence for a strong causal link.

**Figure 2.**
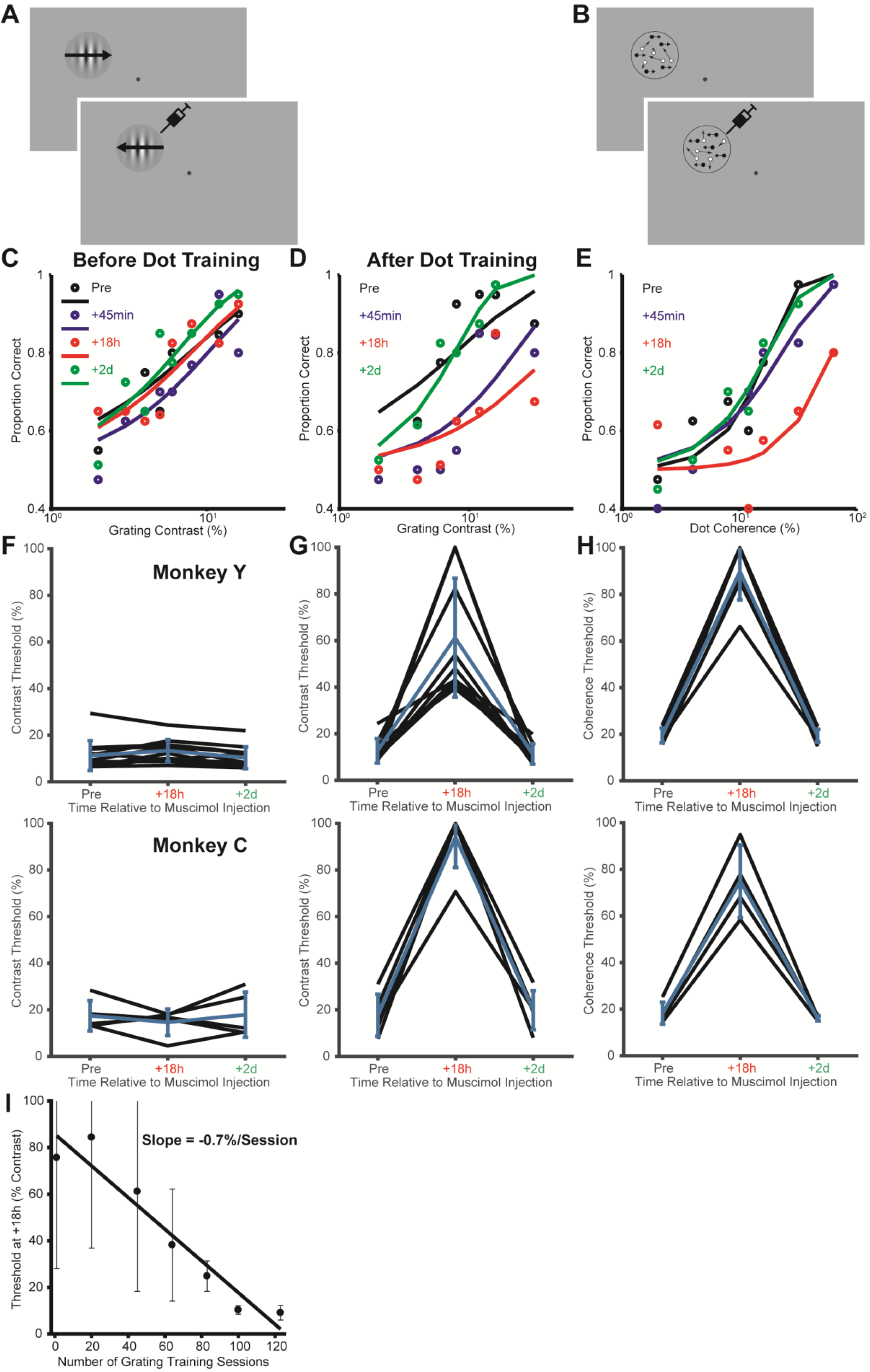
Training with random-dot stimuli alters the causal contribution of MT to grating motion direction discrimination. **(A)** Schematic illustration of the grating motion direction discrimination task. **(B)** Schematic illustration of random-dot motion direction discrimination task. **(C)** Data from a representative experiment *before* training on the random dot motion direction discrimination task. 40 trials were performed at each stimulus condition. The lines indicate Weibull functions fit to the data at different time points following inactivation. The fit was obtained with the maximal likelihood procedure. **(D and E)** Data from a representative experiment *after* training on the random dot motion direction discrimination task. The lines indicate Weibull functions fit to the data at different time points following inactivation. **(F)** Summary of inactivation effects *before* training on the random dot motion direction discrimination task. The figure shows the psychophysical threshold at different time points following inactivation. The blue line is the mean ± std. **(G and H)** Summary of inactivation effects *after* training on the random dot motion direction discrimination task. The data shows the psychophysical threshold at different time points following inactivation. **(I)** Plasticity induced by dot training is reversible and dependent on training. Each data point indicates the psychophysical threshold at +18h after injection. The threshold is plotted against the number of grating training sessions after the dot training had stabilized. Error bars indicate 95% confidence interval of the threshold estimates. The line indicates a linear regression fit with slope of -0.7%/session.

## Contribution of MT to motion perception following training with random dot stimuli

We next trained the same animals to perform the same motion discrimination task, but with random dots as the stimulus. In this case, we manipulated task difficulty by changing dot coherence (Britten et al., 1992), and again found that performance improved over the course of several weeks (**Figure 1G**). For both animals, performance saturated at a coherence threshold of around 15%. We then continued with neurophysiological recordings.

For the random-dot stimulus, we again found that CP values were on average slightly but significantly above chance (0.523 ± 0.060, permutation test; *P* < 0.001). The median of the CP distribution for dots was not significantly different from that found with gratings (permutation test; *P* = 0.31). However, in contrast to the results of the grating experiment, we found that MT inactivation severely impaired behavioral performance (**Figure 2E** **and** **H**), leading to an average increase in psychophysical threshold of 65 ± 13% (70 ± 12% for monkey Y and 57 ± 11% for monkey C), when animals were tested 18 hours after the injection. This is consistent with previous work indicating a strong, causal role for MT in motion discrimination for random dot stimuli (Chowdhury and DeAngelis, 2008; Newsome and Pare, 1988; Rudolph and Pasternak, 1999).

Following the training and testing with random dot stimuli, we repeated the grating experiments in the same animals. Overall behavioral performance on this task remained high, with no significant change from that observed before training on the random dots stimulus (**Figure S5C, D**; Wilcoxon rank sum test; *P* = 0.94). However, the effect of MT inactivation was dramatically different: After training with random dot stimuli, muscimol injections led to a very large decrease in performance on the grating motion task (**Figure 2D** **and** **G**): The average threshold increase at 18 hours after the injection was 59 ± 27% (49 ± 27% for monkey Y and 76 ± 15% for monkey C) (Wilcoxon rank sum test compared to the threshold change before dot training; *P* < 0.001; *P* < 0.001 for monkey Y and *P* = 0.008 for monkey C; **Figure 2** **and** **S6**). Smaller changes in threshold were evident for both stimuli when the animals were tested at 45 minutes after muscimol injections, with performance returning to baseline 48 hours later (**Figure S6**). Thus training on the random-dot stimulus led to a greater role for MT in perceptual decisions about motion, even for grating stimuli, which elicit strong direction selectivity in other areas. This increased role for MT appears to be obligatory, since the results shown in Figure 2C indicate that, during muscimol injections, the animals could have achieved far better task performance by relying on areas other than MT.

To ensure that the increase in psychophysical threshold after muscimol injection was not due to a direct effect of injection pressure or to a loss of attention or motivation during the experiments, we tested the animals with stimuli placed in the hemifield ipsilateral to the injection site. In this case, psychophysical thresholds were unaffected (**Figure S2**).

To further examine the plasticity of the perceptual readout, we carried out a third round of training in one animal: Following the random dot experiments, we again trained the animal repeatedly on the grating discrimination task, with no further exposure to the random dots. In this case the role of MT in perceptual decisions, as assessed with muscimol injections into the same part of MT (**Figure S4**), declined steadily over the course of many sessions (**Figure 2I**). A linear regression on the threshold at 18h post muscimol injection versus training session showed a significant decrease (*F* test that the slope is zero; *P* < 0.001), with the slope of the best-fitting line indicating that the effect of MT inactivation on perceptual threshold decreased by 0.7% per session. This is not likely to be an effect of repeated muscimol inactivation, since permanent lesion experiments often show much faster recovery (Yamasaki and Wurtz, 1991).Thus the contribution of MT to perceptual decision-making could be decreased or increased with training.

## Comparison of muscimol effects before and after random dot training

To ensure that these results were not due to differential administration of muscimol at different time points during the experiment, we carried out a number of control analyses. In particular, we verified that the injection sites were the same before and after dot training, based on both the physical position of the electrodes and the receptive field position mapped prior to injection (**Figure S1A, B**). Overall there were no significant differences in the receptive field eccentricities mapped before and after dots training in monkey Y (6.6 ± 1.7° before training and 8.0 ± 1.8°; Wilcoxon rank sum test, *P* = 0.12) or in monkey C (7.7 ± 4.1° and 7.1 ± 4.1°; Wilcoxon rank sum test, *P* = 0.99). We also verified that activity was consistently abolished following muscimol injection, with spike rates going to zero in all sessions (**Figure S1C and D**), and the evoked local field potential amplitude decreasing by about 90% following muscimol inections (**Figure S1E**). These effects were similar before and after dot training (**Figure S1E**; Wilcoxon rank sum test, Monkey Y: *P* = 0.81; Monkey C: *P* = 0.24). These data indicate that the same quantity of muscimol was injected into the same MT sites across all phases of the experiments in both animals.

## Impact of training on single-neuron correlations with behavior

Following training with random dots, the increased contribution of MT to motion perception could reflect an increase in the motion sensitivity of individual MT neurons, or it could reflect an increased weighting of MT outputs by downstream cortical areas (Law and Gold, 2008, 2009). To examine these possibilities, we quantified the stimulus sensitivity of individual MT neurons by computing the Receiver Operating Characteristic (ROC) score for each stimulus condition (**Figure 3A**, **B** **and** **C**) (Green and Swets, 1966). The ROC score captures the fidelity with which the firing rate of a neuron encodes a stimulus. Moreover, the correlation of a neuron’s ROC score with its CP value can reflect the selectivity of the readout from an area (Law and Gold, 2008; Masse et al., 2012). We compared these quantities for MT responses to the same grating stimuli, using neurons recorded before (n = 79; n = 49 for Monkey Y and n = 30 for Monkey C) and after (n = 55; n = 42 for Monkey Y and n = 13 for Monkey C) training with random dot stimuli.

**Figure 3.**
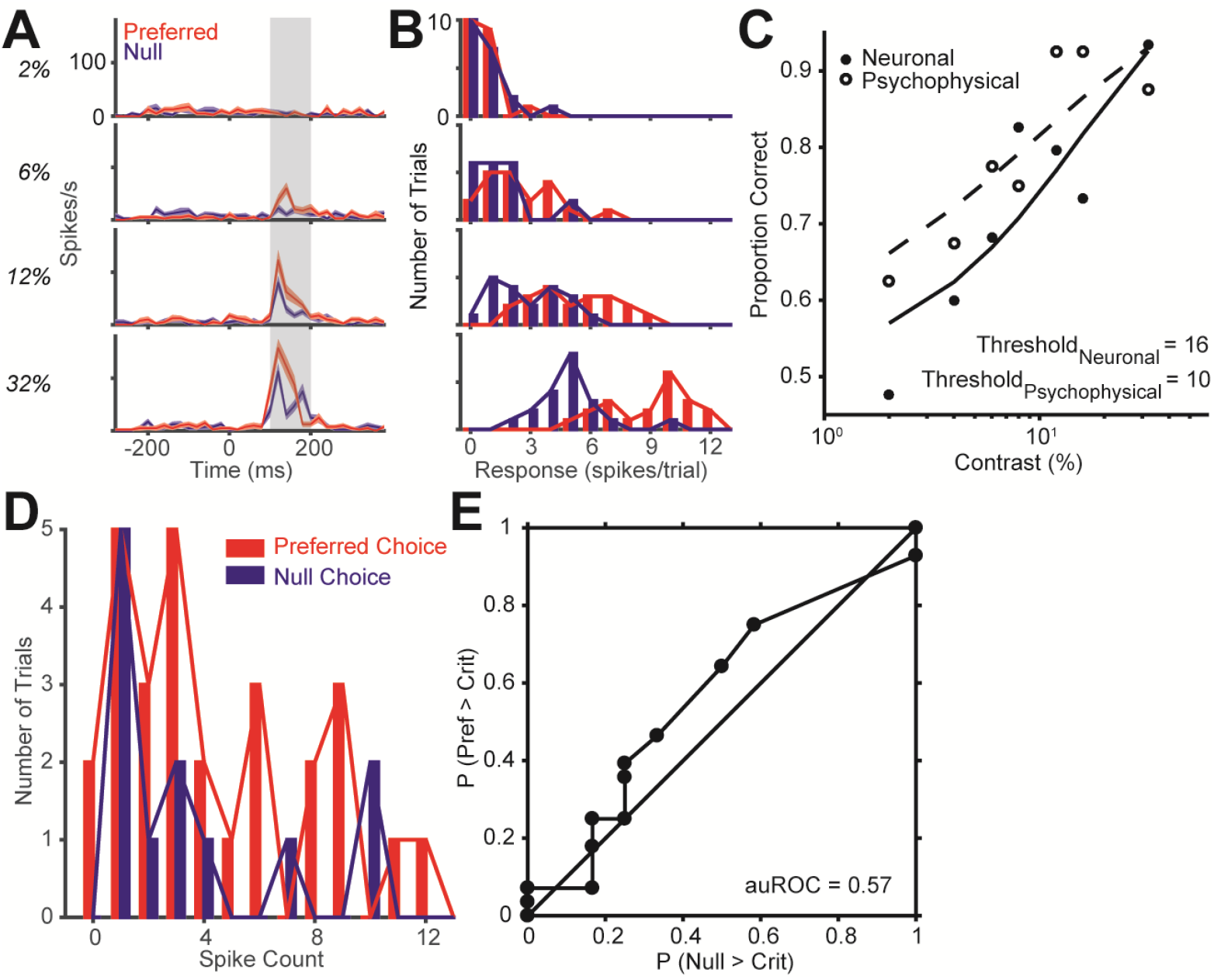
Quantification of neurometric performance and correlation between single-neuron activity and perceptual decisions. (**A**) Peri-stimulus time histogram (PSTH) of the responses for an example neuron. Red represents the responses to the preferred direction, and blue represents responses to the null direction. (**B**) Distribution of firing rate between 100 and 200 ms after stimulus onset. (**C**) The neurometric function computed as the area underneath the ROC curves for each stimulus contrast (solid symbols fitted by the solid line). The corresponding psychometric function on that session is superimposed (open symbols fitted the dashed line). The 92% correct thresholds are indicated below. (**D**) The distribution of firing rates between 100 and 200 ms for a grating stimulus at 0% contrast, grouped according to the choice the animal made. (**E**) The ROC curve for the distributions yields an area underneath the ROC score of 0.57. The reported CP for each neuron was computed by averaging the CP across all stimulus conditions, for each condition in which the monkey made at least five choices for each direction.

We found that training with random dots had no consistent effect on ROC scores across the MT population (Wilcoxon rank sum test; *P* = 0.64; *P* = 0.60 for Monkey Y and *P* = 0.59 for Monkey C), indicating that training did not change the sensitivity of individual neurons to the encoding of the stimulus; there was also no significant change in CP for the same population (Wilcoxon rank sum test; *P* = 0.49; *P* = 0.77 for Monkey Y and *P* = 0.55 for Monkey C). However, following training on random dot stimuli, CP values became more strongly correlated with the sensitivity of individual neurons, as captured by the neurometric thresholds (Britten et al., 1992). This relationship was weak and marginally significant before training with random dot stimuli (**Figure 4A**, *r* = -0.20, *P* = 0.07; *r* = -0.16, *P* = 0.28 for Monkey Y and *r* = -0.28, *P* = 0.14 for Monkey C), but stronger following training (**Figure 4B**, *r* = -0.32, *P* = 0.02; *r* = -0.30, *P* = 0.05 for Monkey Y and *r* = -0.40, *P* = 0.18 for Monkey C). A bootstrap analysis (see Methods) yielded a stronger, negative dependence of CP on neurometric threshold after dots training (Wilcoxon rank sum test; *P* < 0.001). Overall, these results are consistent with the idea that training with random dots does not improve the sensitivity for such stimuli in MT, but leads to a more selective readout of MT neurons by downstream decision-making areas (Law and Gold, 2008, 2009).

**Figure 4.**
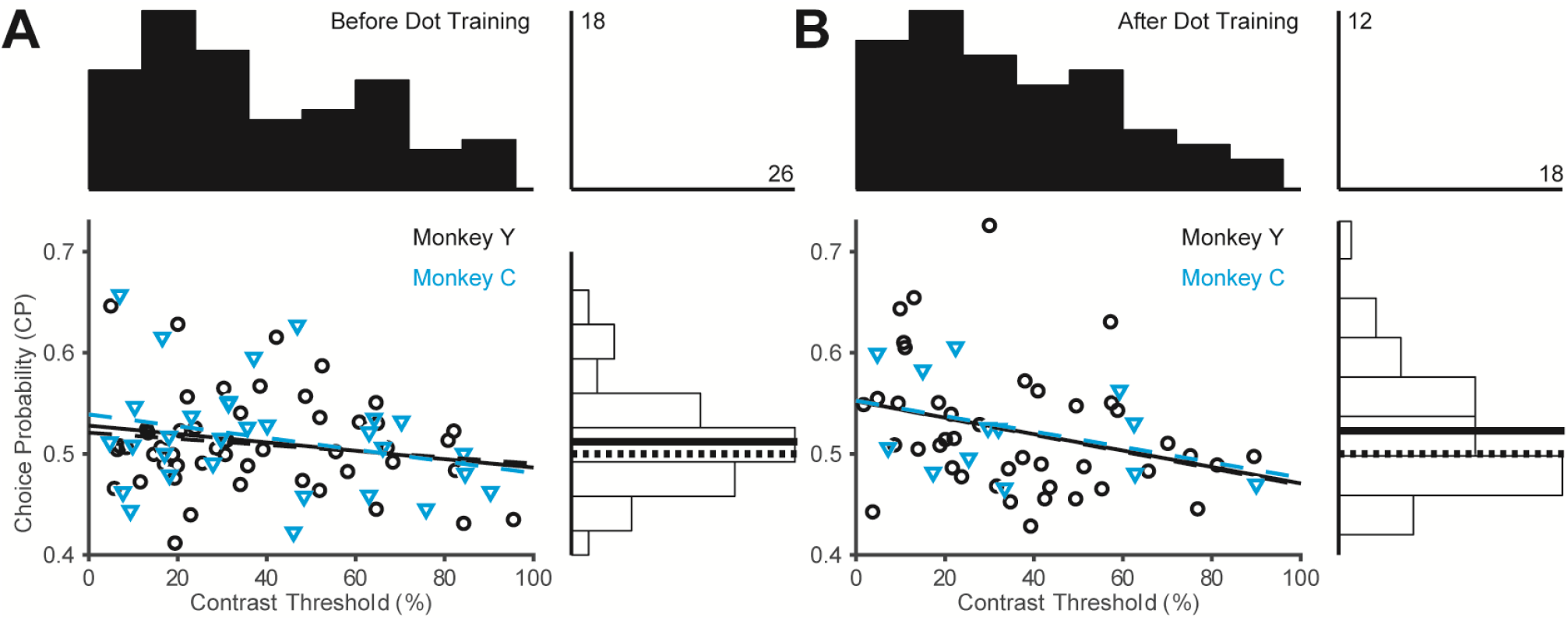
The relationship between choice probability (CP) and neurometric thresholds before and after dot training. **(A)** Scatter plot of CP and neurometric thresholds before dot training. The black circles indicate the data from Monkey Y and the blue triangles indicate the data from Monkey C. The lines indicate linear fits. The black dashed line indicates the linear fit for Monkey Y, and the blue dashed line indicates the fit for Monkey C. The marginal distributions of CP and neurometric thresholds are also shown. The solid line indicates the median CP value, and the dotted line is at 0.5. The scale bars indicate the number of neurons in the highest bin. **(B)** CP and neurometric thresholds for the same grating task after dot training.

## Influence of MT microstimulation depends on training history

The single-neuron results (**Figure 4**) suggest that training did not improve the sensitivity of the neurons, but training might have caused the decision process to become more correlated with activity in the most sensitive MT neurons. To test whether this correlation had a causal dimension, we performed microstimulation experiments only in monkey Y before and after random dots training. We used stimulation parameters (see Methods) that influence neural activity on a scale comparable to that of a cortical column (Murasugi et al., 1993; Salzman et al., 1992).

Figure 5A shows an example psychometric function for grating motion discrimination prior to dot training. Compared to non-stimulated trials (dashed line), microstimulation led to a pronounced decrease in the slope of function, which was consistent across recording sites (**Figure 5G**; mean change -0.015 ± 0.023; Wilcoxon signed rank test; *P* = 0.003). This suggests that MT contributed substantial noise to the decision making process, as would be expected if the weighting of MT outputs was independent of the sensitivity of the neurons (**Figure 4** **A**). Consistent with this idea, the microstimulation-induced bias toward the preferred directions of the stimulated sites was weak and non-significant (**Figure 5A, D and G,** 0.7 ± 0.7 for bias; Wilcoxon signed rank test; *P* = 0.19).

**Figure 5.**
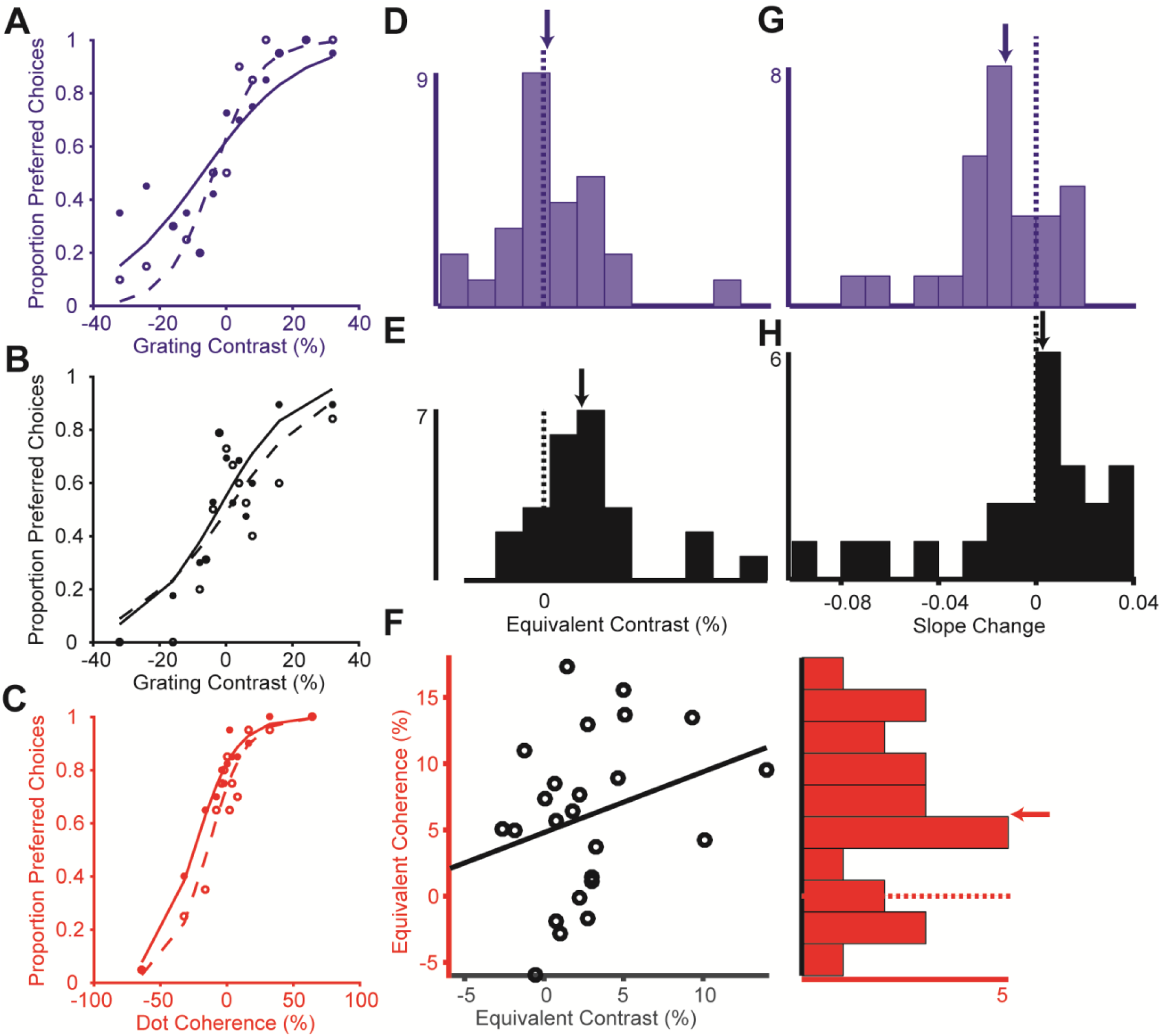
Effects of microstimulation before and after dot training in monkey Y. **(A)** Psychophysical grating performance for an example experiment before dot training (blue). The lines indicate logistic functions fit to the data. Each point is based on 20 trials. The non-stimulated trials are represented by the open circles and fitted with the dashed lines. The stimulated trials are represented by the filled circles and fitter with the solid lines. The same convention is used throughout the figure. **(B)** Psychophysical performance on the grating task for an example experiment after dot training (black). **(C)** Psychophysical performance on the random dots task for an example experiment (red). **(D)** Direction bias induced by microstimulation for the grating experiments before dot training. Scale bar indicates the number of stimulation sites in the highest bin. The arrow indicates the median of the population. The dotted line indicates zero. **(E)** Microstimulation-induced bias for the grating experiments after dot training. **(F)** Scatter plot of the microstimulation bias for dot experiments against the bias for the grating experiments. The marginal distribution on the right is the bias for the dot experiments. **(G)** Change in the slope of the psychometric function induced by microstimulation for the grating experiments before dot training. **(H)** Slope change for the grating experiments after dot training.

After training with the random-dot stimulus, the bias induced by microstimulation at the same sites (**Figure S7**) increased significantly, and the effect on the slope of the psychometric function was smaller (**Figure 5B, E and H,** 2.8 ± 0.8 for bias and 0.021 ± 0.049 for slope change, Wilcoxon rank sum test compared to before dot training; *P* = 0.040 and 0.047, respectively). Thus training with dots caused the predominant psychometric effect of microstimulation to change from a reduction in slope to a horizontal shift (bias), despite the fact that stimulation parameters were identical in both experiments.

As in previous studies, we found that this bias was directed toward the preferred direction of the stimulation sites for the dot stimulus (Wilcoxon signed rank test; *P* < 0.001) (Salzman et al., 1990; Salzman et al., 1992) (**Figure 5C and F**). Interestingly, this bias for random dot stimuli was correlated with the bias for grating stimuli at the same site (**Figure 5F**, *r* = 0.29, *P* = 0.05). Thus an additional effect of training appears to be the development of representations of motion that are invariant to the spatial structure of the stimulus.

## Constraints on the cortical sites for grating motion discrimination

The results thus far suggest that MT has little causal influence on motion perception following training with drifting gratings. A reasonable assumption is that training with gratings leads to a greater dependence on lower-level areas, such as V1, V2, or V3, for perceptual decisions about motion. To test this idea, we examined behavioral performance on a task involving high-contrast gratings of different sizes (**Figure 6A**). This provides a measure of spatial integration (Liu et al., 2016), which is expected to be weaker in low-level areas, where neurons have relatively small receptive fields (Angelucci and Bressloff, 2006).

**Figure 6.**
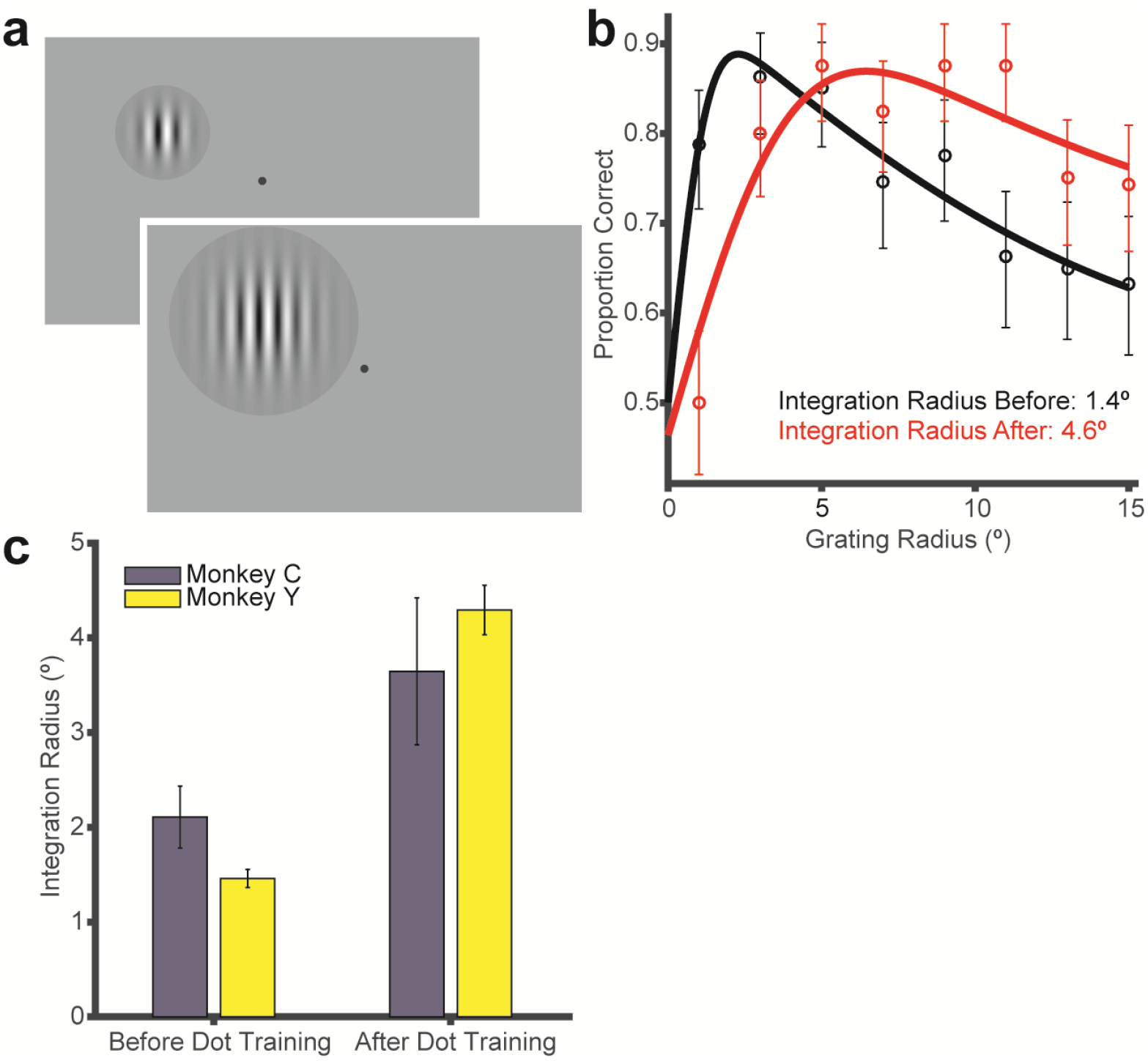
The range of psychophysical spatial integration is larger after training with random dot stimuli. **(A)** Schematic illustration of the paradigm in which grating size in visual space was varied. **(B)** Example psychometric functions before (black) and after (red) training with random-dot stimuli. The lines indicate the fit of a difference of error functions to the data. Error bars indicate 95% binomial proportion confidence interval. **(C)** Psychophysical integration radius for the 2 monkeys before and after training with random dots; mean ± s.e.m.

Consistent with previous work (Liu et al., 2016; Tadin et al., 2003), we found that performance increased with stimulus size up to a certain point (**Figure 6B**), beyond which it declined, presumably as a consequence of surround suppression (Liu et al., 2016; Tadin et al., 2003; Tadin et al., 2011). For these behavioral effects we estimated the animals’ psychophysical integration radius (see Methods) and found that it was larger after random-dot training (red) than before (black), as would be expected if training with random-dot stimuli shifted perceptual processing to a region with larger receptive fields. The mean integration radius was 1.9 ± 1.1o before training with random dot stimuli (**Figure 6C**) and 4.0 ± 1.8o afterwards (**Figure 6C**, Wilcoxon rank sum test, *P* = 0.02).

Finally, we examined the retinotopic specificity of the observed training effects. Following training on random dot stimuli placed in one visual hemifield, we tested behavioral performance on drifting grating stimuli placed in the opposite hemifield (**Figure 7A**), as in **Figure S2**. However, in this experiment we injected muscimol into MT in the cortical hemisphere opposite the trained one. In this case, there was little effect of muscimol on perceptual performance (**Figure 7B** **and** **C**), even though the activation targeted the area of MT that represented the stimulus. In this regard the effects of muscimol were similar to those reported previously before training with random-dot stimuli (**Figure 2F**). This suggests that the effect of training with random-dot stimuli was specific to the trained hemifield.

**Figure 7.**
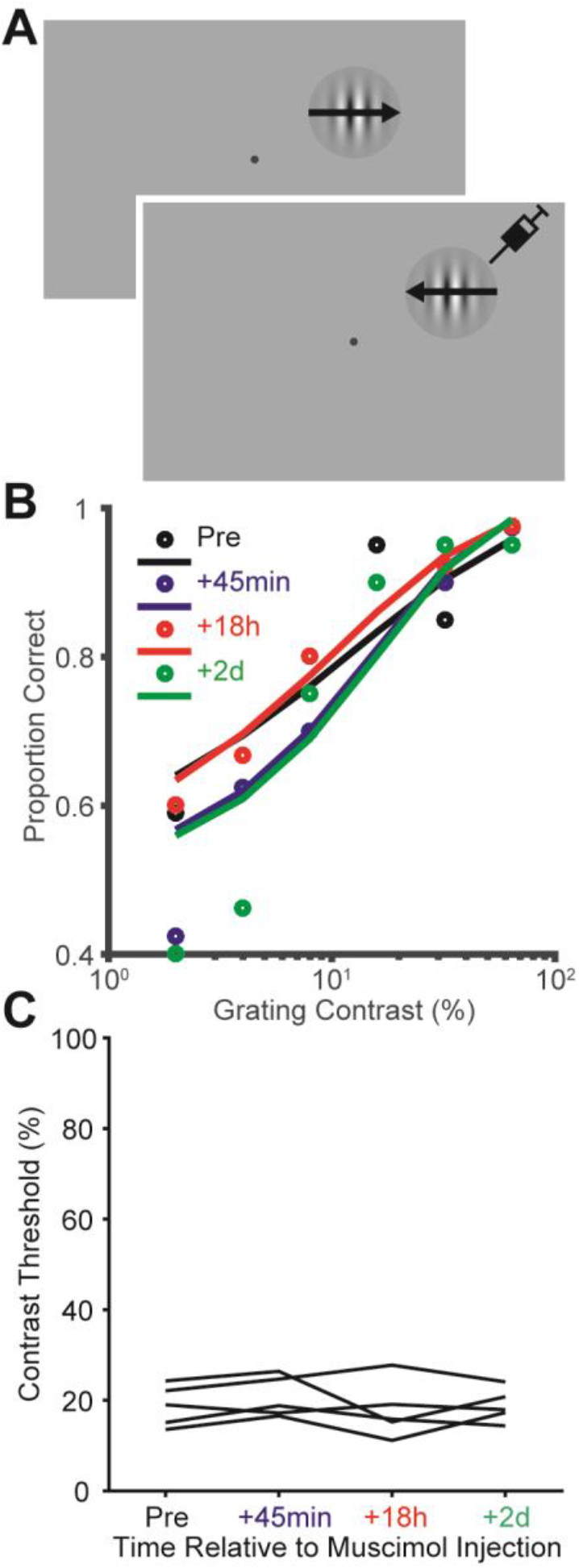
Spatial specificity of the effects of training with random dots. These experiments were conducted after training with stimuli in the *left* visual hemifield. **(A)** Schematic illustration of the grating motion direction discrimination task with the stimulus placed in the *right* visual hemifield. **(B)** Data from a representative experiment with drifting gratings, after training on random-dot stimuli in the left visual hemifield. The lines indicate Weibull functions fit. The figure shows little change in psychophysical performance at each time point following inactivation. **(C)** Summary of inactivation effects on grating motion perception, after random-dot training. The data shows the psychophysical threshold at different time points following inactivation.

## Discussion

We have shown that area MT, the brain region most clearly linked to visual motion processing, is not necessarily involved in visual motion perception. Rather, the causal link depends heavily on recent perceptual experience: For drifting grating stimuli, manipulation of MT activity can dramatically alter motion perception, or it can have little effect, depending on recent training history. Additional analyses show that the readout of visual motion information can shift flexibly between lower-level and higher-level cortical areas.

### Relationship to previous literature

Area MT is also involved in processing depth information from retinal disparity cues (Maunsell and Van Essen, 1983), so that perceptual decisions about disparity also rely on MT activity (DeAngelis et al., 1998). A previous study showed that the causal relationship between MT and disparity perception can be altered with training (Chowdhury and DeAngelis, 2008). Our results showed that the converse is also possible: Training on a motion discrimination task that specifically recruits MT can strengthen its contribution to other motion perception tasks.

Within the domain of motion processing, a previous imaging study also showed that the contribution of MT depends on training. The authors trained human subjects with a 100% coherent dot pattern, and showed that such training can actually reduce the causal contribution of MT to motion perception (Chen et al., 2016). These results might seem to contradict ours, but it is important to consider that in our study, the monkeys were trained with low-coherence dot patterns. This is a crucial difference, as it encourages the subjects to integrate motion information over space, favoring cortical areas with larger receptive fields (**Figure 6**). In contrast, the 100% coherent dot field contains local information about motion at every point in the stimulus, and in this regard it is more analogous to our drifting grating stimulus. Thus the two sets of results can be reconciled under the idea that training with stimuli that contain informative local cues lead to reliance on lower-level cortical areas, while noisy stimuli are processed most effectively by higher areas such as MT (Rudolph and Pasternak, 1999).

The study by Chen et al. (2016) also reported that the encoding of motion in MT, as measured with fMRI, changed with training. We did not find evidence for such a change at the single-neuron level (**Figures 3** **and** **4**); rather our recording and stimulation results (**Figure 5**) are more consistent with the idea that training changes the read-out of the MT population by downstream areas (Law and Gold, 2008). A similar conclusion was reached in a study that used microstimulation to probe the transmission of signals from MT to downstream areas (Seidemann et al., 1998).

More generally, our results add to previous observations that cortical activity can be correlated with behavior in the absence of any causal relationship (Katz et al., 2016; Nienborg and Cumming, 2009; Zenon and Krauzlis, 2012). Although previous studies of MT and motion perception in monkeys have hinted at possible training effects (Chowdhury and DeAngelis, 2008; DeAngelis and Newsome, 2004; Liu and Newsome, 2005), our results suggest that this factor is critical in interpreting correlations between neurons and behavior. Similarly, human studies frequently make use of grating stimuli to infer a connection between behavioral performance and the properties of MT, without necessarily considering the role of training, e.g., (Churan et al., 2009; Tadin et al., 2003; Yo and Demer, 1992). Future research could detect a potential influence of training by using non-invasive inactivation to perturb cortical activity (Chang et al., 2014; Walsh et al., 1998).

Previous studies on sensory decision-making have considered the flow of neural signals from brain regions that represent sensory information to those that represent decisions (Gold and Shadlen, 2007; Mazurek et al., 2003; Shadlen et al., 1996). A common conception is that the sensory transformation is hierarchical, with early stages being less involved in perceptual decisions (Nienborg and Cumming, 2006; Tsunada et al., 2011). This may be the default state of system, as the invariance of such representations to irrelevant stimulus dimensions simplifies decision-making (DiCarlo and Cox, 2007). However, the evidence on this point is scant and possibly confounded by feedback influences (Haefner et al., 2013; Nienborg and Cumming, 2009).

In the auditory domain, there appears to be a feedforward pathway from the middle-lateral to the anterolateral cortex, with the latter being more closely tied to perceptual decisions (Tsunada et al., 2016). However, both areas receive input from primary auditory cortex, and the relative influences of these areas to perceptual decisions among these areas are likely to be task-dependent (Niwa et al., 2013). In the somatosensory domain, there is some evidence that the second somatosensory area has a stronger influence on decisions than primary somatosensory cortex (Romo et al., 2002). However, as in the auditory domain this relationship is not entirely straightforward. It would be interesting to examine the effects of training in these other domains.

### Implications for perceptual learning

Perceptual learning can be associated with an improvement in the sensitivity of individual neurons in early sensory areas (Adab and Vogels, 2011; Raiguel et al., 2006; Schoups et al., 2001), or to plasticity in the decision areas that read out the sensory evidence (Law and Gold, 2008). Consistent with the latter idea, we found that training with random-dot stimuli led primarily to a change in the apparent weighting of neurons according to their sensitivity (**Figure 4**), with no improvement in performance with grating stimuli (**Figure S5C, D**).

In the psychophysics literature, perceptual learning studies are often concerned with the transfer of learning across stimulus dimensions, particularly spatial position (Hung and Seitz, 2014; Sagi, 2011; Xiao et al., 2008). Our results are consistent with previous psychophysical studies indicating the spatial specificity of training effects (Ahissar and Hochstein, 1996, 1997; Karni and Sagi, 1991) (**Figure 7**). Less psychophysical work has been devoted to the question of generalization across stimulus cues, and the results have been somewhat contradictory (Ivanchenko and Jacobs, 2007; Pilly et al., 2010; Rivest et al., 1997). Our results suggest that the neural readout of a single stimulus quantity (motion direction) depends strongly on the subject’s experience with the particular cue that defines the quantity. However, different training procedures can produce different levels of specificity in perceptual learning (Das et al., 2012; Hung and Seitz, 2014; Wang et al., 2016; Xiao et al., 2008), and these differences can be explained based on changes in the relative weighting of low and mid-level cortical areas (Dosher et al., 2013; Talluri et al., 2015). It will be important to examine the influence of different training regimens in future animal studies.

Our findings can also be interpreted within the framework of reverse hierarchy theory (Ahissar and Hochstein, 2004; Hochstein and Ahissar, 2002), which posits that learning of difficult stimulus discriminations leads to a greater reliance on lower-level cortical areas. In this framework, the default state of the visual system would be to base perceptual decisions about motion on activity in higher-level cortical areas such as MT or MST, which represent global aspects of motion stimuli (Born and Bradley, 2005; Khawaja et al., 2013; Mineault et al., 2012). Training on a difficult grating discrimination task would shift the readout toward lower-level cortical areas that represent the local motion of the stimulus. Our results on the behavioral integration radius for grating stimuli (**Figure 6**) are entirely consistent with this idea.

## Author Contributions

L.D.L. and C.C.P. conceived and designed this study. L.D.L. performed the experiments and analyzed the data. L.D.L. and C.C.P. wrote the paper.

## Acknowledgements

This work was supported by grants from the Canadian Institutes of Health Research to C.C.P. (PJT-148488) and to L.D.L. (CGSD-121719). We would like to thank Julie Coursol and the staff of the Animal Care Facility (Montreal Neurological Institute) for excellent technical support, and A. Seitz and R. Born for helpful discussions.

## STAR Methods

### Key Resources Table

| REAGENT or | SOURCE | IDENTIFIER |
| --- | --- | --- |
| Experimental Models: Organisms/Strains |  |  |
| Rhesus macaque<br>(*Mucaca mulatta*) | N/A | N/A |
| Software and Algorithms |  |  |
| MATLAB | MathWorks | https://www.mathworks.com/products/matlab.html |
| Plexon Sort Client | Plexon | http://www.plexon.com/products/software |
| Plexon Offline Sorter | Plexon | http://www.plexon.com/products/offline-sorter |
| Other |  |  |
| Plexon MAP System | Plexon | http://www.plexon.com/products/map-data-acquisition-system-plexon |
| Plexon V probes | Plexon | http://www.plexon.com/products/plexon-v-probe |
| Muscimol | Sigma | Sigma M1523-25MG |
| Blackrock CereStim 96 | Blackrock | http://blackrockmicro.com/neuroscience-research-products/ephys-stimulation-systems/cerestim-96-neurostimulation-system/ |
| Microelectrodes | FHC | FHC UEPLECSEBN1E |

### Contact for Reagent and Resource Sharing

Further information and requests for resources and reagents should be directed to and will be fulfilled by the Lead Contact, Christopher C. Pack.

### Experimental Model and Subject Details

Two adult female rhesus monkeys (*Macaca mulatta*) (Monkey Y, age: 10 years; Monkey C, age: 8 years, weight: both 7 kg) participated in this study. Before training, under general anesthesia, an MRI-compatible titanium head post was attached to each monkey’s skull. The head posts served to stabilize their heads during subsequent training and experimental sessions. For both monkeys, eye movements were monitored with an infrared eye tracking system (EyeLink1000, SR Research) with a sampling rate of 1,000 Hz. We chronically cemented the recording cylinders to the monkeys’ skulls to access MT from a dorsal-posterior approach. The recording cylinders were placed 16 mm lateral to the midline and 16 mm dorsal to the occipital ridge of the skulls. The angle of approach in the parasagittal plane was 20° above the horizontal.

All procedures conformed to the regulations established by the Canadian Council on Animal Care and were approved by the Institutional Animal Care Committee of the Montreal Neurological Institute.

## Method Details

*Electrophysiological recordings, pharmacological injections, and microstimulation* Area MT was identified based on an anatomical MRI scan, as well as depth, prevalence of direction-selective neurons, receptive field size to eccentricity relationship, and white matter to grey matter transition from a dorsal-posterior approach. We used a system in which a grid was rigidly positioned within the recording cylinder. We then lowered the electrodes every day through the same grid holes to the same depth (**Figure S1B**). We recorded single units using linear microelectrode arrays (V-Probe, Plexon) with 16 contacts.

At the beginning of each experiment, we lowered the electrode array to the appropriate depth and then estimated multi-channel receptive fields by manually positioning a moving bar in the visual field (**Figure S1A and B**). Visual motion stimuli were displayed at 60 Hz at 1,280 by 800 pixels resolution; the viewing area subtended 60° × 40° at a viewing distance of 50 cm. The neuronal signals were monitored continuously during acquisition by computer display, and the spike signals were band-pass filtered at 150 Hz-8 kHz and monitored on an oscilloscope and loudspeaker. The spike signals were thresholded online, and spikes were assigned to single units by a template-matching algorithm (Plexon MAP System). Offline, spikes were manually sorted using a combination of automated template matching, visual inspection of waveforms, clustering in the space defined by the principle components, and absolute refractory period (1 ms) violations (Plexon Offline Sorter).

Direction and speed preferences were quantified using 100% coherent dot patches placed inside the receptive fields. Offline, the receptive field locations were further quantified by fitting a spatial Gaussian to the neuronal response measured over a 5 x 5 grid of stimulus positions. The grid consisted of moving dot patches centered on the initially hand-mapped receptive field locations. We confirmed that all neurons included in our analysis had receptive field centers within the stimulus patch used for the behavioral experiments.

*Injection of muscimol*. The linear array contained a glass capillary with an inner diameter of 40 μm. One end of the capillary was located between contacts 5 and 6 of the array (contact 1 was most dorsal-posterior), and the other end was connected via plastic tubing to a Hamilton syringe. Muscimol (typically 2 μL at 0.05 μL/min) was injected with a mini-pump. The concentration of muscimol was 10 mg/mL (Chowdhury and DeAngelis, 2008). We verified that neural activity had ceased before commencing behavioral experiments at 45 minutes after injection (**Figure S1C, D and E**).The injection of muscimol and the initial behavioral testing were performed in the afternoon of the first day, and behavioral testing was repeated at intervals of 18 hours and 2 days. We also verified that the behavioral deficit was localized within the vicinity of the RF location of the neuron near the injection site. We performed perimetry to test the size of the scotoma for a typical injection (**Figure S3**). Experiments with muscimol were conducted no more than once per week.

*Microstimulation*. For the microstimulation experiments, we used single-contact platinum-iridium electrodes (∼0.5 MΩ at 1 kHz, FHC). The electrodes were lowered into area MT until single units were well isolated online. Stimulation was delivered using an isolated constant-current stimulator (CereStim 96, Blackrock Microsystems) with symmetric, biphasic pulses, 200 μs in total duration (cathodal followed by anodal, 100 μs each). We used a stimulation frequency of 200 Hz and current of 10 μA, since previous studies found that these parameters can produce robust behavioral effects (Murasugi et al., 1993; Salzman et al., 1990). Stimulation was delivered from 100 ms to 200 ms after the onset of the visual stimulus, corresponding to the time at which we found robust directional selective responses (**Figure 3A**).

### Training

Animals were trained to perform a two-alternative, forced choice, coarse motion direction discrimination task, using operant conditioning. The initial instruction stimulus was a 5° radius, high contrast vertical grating placed at 5° eccentricity. Both monkeys initially exhibited a lateral bias towards one of the targets. This stereotypical behavior was eliminated by inserting “correction” trials after three consecutive guesses towards the biased target. Correction trials are trials with the stimulus moving in the direction opposite that of the behavioral bias. The correction trials were removed after the elimination of the bias, which gradually subsided over one month of training. We began tracking the animals’ performance when performance at the highest contrast reached 75% correct (**Figure 1D**). The psychophysical contrast threshold for the grating motion direction discrimination saturated after approximately one month of training (**Figure 1F**).

Following this initial training period, we further generalized the discrimination of grating motion for eight different directions, different spatial and temporal frequencies, and various spatial locations. This generalization required an additional three months. We only started our experiments after the psychophysical thresholds appeared to saturate for the stimulus locations in our study (**Figure 1F,** **S5**). After training on the grating motion discrimination task, the task with random dots required much less time to train and generalize. The animals’ performance at high stimulus coherences were already above 75% correct in the first session of training (**Figure 1E**). The psychophysical coherence threshold saturated in approximately the same number of sessions as for the grating motion discrimination (**Figure 1G**).

### Motion direction discrimination tasks

The structure of an individual trial is illustrated in **Figure 1A**. Each trial began with the onset of a fixation point. The monkey was required to establish and maintain fixation within a 2° × 2° window for 300 ms, after which a drifting Gabor or random dots patch appeared on the receptive field centers (Mean eccentricity = 6.6 ± 1.7°). The spatial and temporal frequency of the Gabor or the speed of the dots were chosen to match the preference of the MT neuron near the injection site. The size of the stimulus also matched the RF size of the MT neuron near the injection site (Mean radius = 6.3 ± 1.2°). The contrast of the Gabor patch or the coherence of the random dots pattern was chosen randomly on each trial from among seven or eight values spanning the range of the monkey’s psychophysical threshold. These values were selected based on previous measurements of the psychometric functions with an emphasis around the steepest sections of the psychometric function. In the experiments in which we manipulated the size of the Gabor patches (**Figure 6**), the size was defined by 2 standard deviations of the Gaussian envelope and ranged from 1° to 15° in steps of 2° (Liu et al., 2016). The motion stimulus was brief (typically 67 ms) in duration, after which the monkey was required to maintain fixation for another 300 ms. At the end of each discrimination trial, the fixation point disappeared, two choice targets appeared, and the monkey made a saccade to the corresponding target to report its perceived motion direction (preferred or null relative to the neurons isolated). The monkey was required to indicate its decision within 0.7 s following the onset of the choice targets. Correct choices were rewarded with a drop of liquid. For the trials that contained no motion signal (0% contrast or 0% coherence), rewards were delivered randomly on half of the trials. If fixation was broken at any time during the stimulus, the trial was aborted. In a typical experiment, the monkeys performed 20-40 repetitions of each distinct stimulus.

### Data analysis

The monkeys’ performance as a function of grating contrast or dot coherence was characterized by fitting a Weibull function to the proportion of correct responses using the maximal likelihood algorithm (FitWeibTAFC in MATLAB). The Weibull function is,

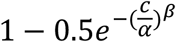

where *α* is the 92% correct threshold; *β* determines the slope of the function; and *c*, is stimulus contrast or coherence.

To quantify the sensitivity of the single neurons, we used the firing rate during the period 100-200 ms after stimulus onset to calculate the area underneath the ROC curve (**Figure 3A and B**). This interval was chosen because the spikes during this time window were significantly correlated with the animals’ behavioral choices (Liu et al., 2016); other time windows between 60-300 ms did not yield results different from those reported here. The auROC scores were plotted as a function of stimulus contrast, and fitted with the Weibull function (**Figure 3C**).

For the microstimulation experiments, psychophysical performance as a function of grating contrast and dot coherence was fitted with a logistic function (**Figure 5**).

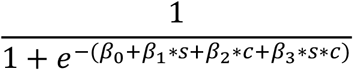

where *β*_0_ is the bias for a particular direction; *β_1_* determines the influence of microstimulation on the function; *s* is 1 for trials with microstimulation and 0 for trials without microstimulation; *β_2_* determines the influence of stimulus contrast or coherence on the function; *c* is stimulus contrast or coherence. *β_3_* determines the interaction between microstimulation and stimulus contrast or coherence. The direction bias injected by microstimulation is reflected by *β_1_*/*β_2_* (**Figure 5D, E and F**). The neural noise injected by microstimulation is reflected by the *β_3_* term (**Figure 5G and H**) (Murasugi et al., 1993).

For the experiments in which we manipulated the size of the Gabor patches (**Figure 6**), psychophysical performance as a function of stimulus size was fitted by the Difference of Error functions (DoE),

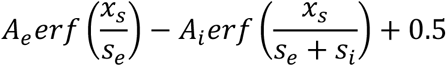

where *A*_*e*_ and *A*_*i*_ scale the height of the excitatory center and inhibitory surround, respectively. *s*_*e*_ and *s*_*i*_ are the excitatory and inhibitory sizes. *x*_*s*_ is the stimulus size (DeAngelis and Uka, 2003; Pack et al., 2005). The integration radii reported in Figure 6 correspond to the parameter *s*_*e*_ for each animal and each training period.

Choice probability (CP) was used to quantify the relationship between behavioral choice and response variability (Britten et al., 1996). For an identical stimulus, the responses can be grouped into two distributions based on whether the monkeys made the choice that corresponds to the neuron’s preferred direction, or the null direction (**Figure 3D**). As long as the monkeys made at least five choices for each direction, ROC values were calculated from these response distributions, and the area underneath the ROC curve gives the CP value (**Figure 3E**). The single CP for each neuron was computed by averaging the CP across all stimulus conditions. The alternative method of z-scoring the data for each stimulus conditions and then combining them into a single pair of distributions for preferred and null choices can underestimate the CP when the number of choices for preferred and null directions differs across stimulus conditions (Kang and Maunsell, 2012).

As we were unable to measure the CP vs. neurometric threshold relationship in the same neurons before and after random dots training, we created matched distributions of neurometric thresholds by subsampling from the original distributions in Figure 4. We first created 5 equal bins of neurometric threshold spanning from 0 to 100, and then resampled randomly to create sub-distributions with equal amounts of data in each bin. We resampled 1,000 times and calculated the slope of the line relating CP to neurometric threshold for each sub-distribution. The distribution of the slopes after training was significantly more negative than before dots training (Wilcoxon rank sum test; *P* < 0.001).

### Quantification and Statistical Analyses

All statistical analyses were performed using built-in MATLAB function and custom scripts. The difference between the contribution of MT before and after dots training was evaluated using the Wilcoxon rank-sum test. We calculated the difference of psychophysical thresholds before and after injection, and we then performed the Wilcoxon rank sum test for the threshold differences. The statistical details regarding the statistical tests used, exact value of n, what n represents, the use of mean or median, and SD or SEM can be found in the main text and/or figure legends. Significance was defined as *P* < 0.05.

## Supplementary Figures

**Figure S1, related to Figure 2.**
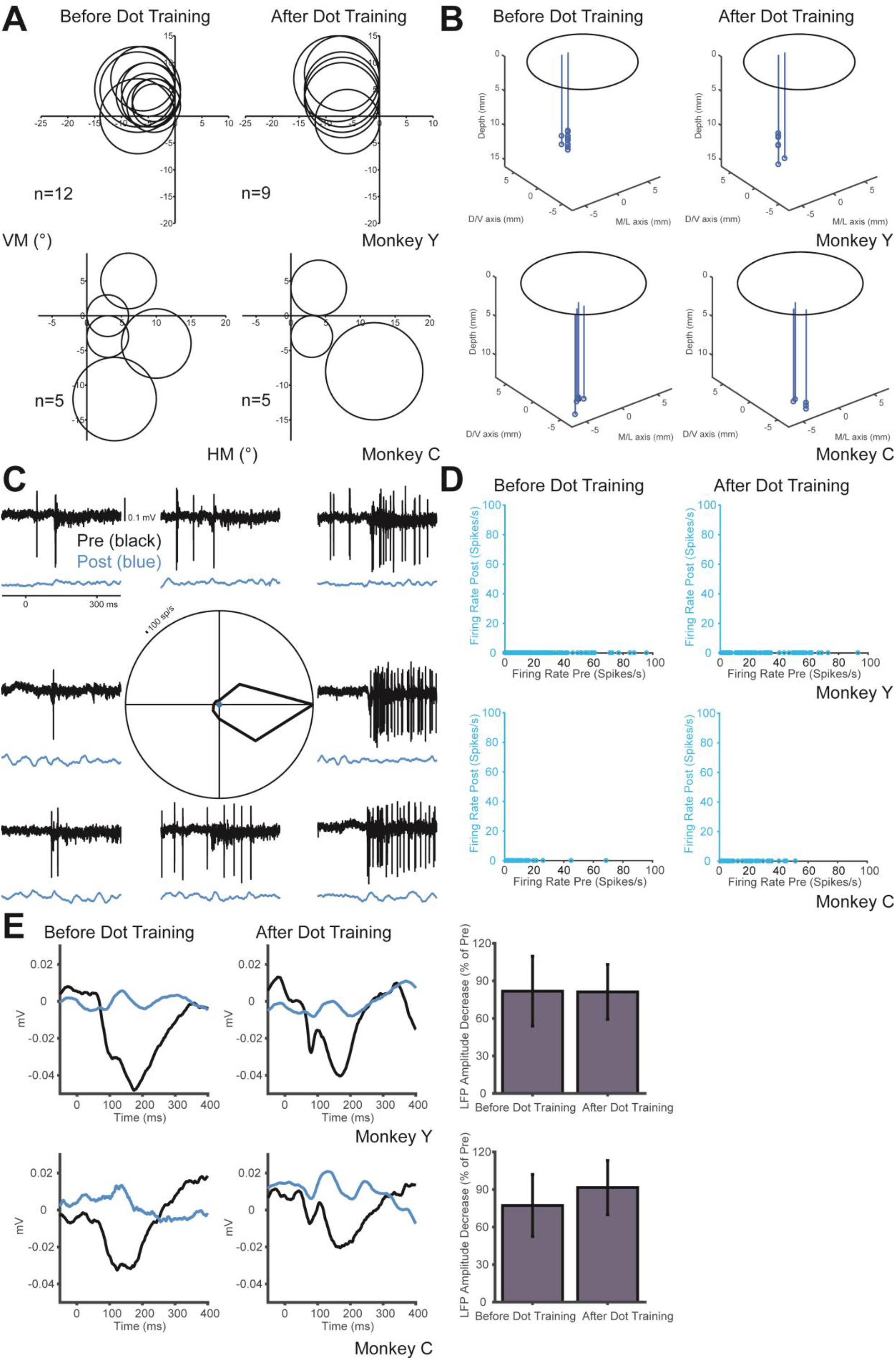
Injections sites were consistent before and after dots training, and the effects of muscimol is robust. **(A)** Receptive field mapping for the neurons near the injection sites. The mean RF eccentricity is 6.6 ± 1.7° before training and 8.0 ± 1.8° after training in Monkey Y (Wilcoxon rank sum test, *P* = 0.12), and 7.7 ± 4.1° and 7.1 ± 4.1° in Monkey C (*P* = 0.99), respectively. The stimulus placement was based on the RF mapping. HM and VM indicate the horizontal and vertical meridian, respectively. **(B)** Reconstruction of the injection sites based on grid positions and depth from the electrode microdrive. The injection sites are indicated by the circles and the injectrode tracks are indicated by the lines. M/L is medial-lateral and D/V is dorsal-ventral. The depth is zeroed at the cortical surface from a dorsal-posterior approach to MT. The mean distance of the injections is 13.0 ± 0.9 mm before training and 13.36 ± 1.6 mm after training in Monkey Y (Wilcoxon rank sum test, *P* = 0.97), and 13.5 ± 0.7 mm and 13.5 ± 0.4 mm in Monkey C (*P* = 0.42), respectively. **(C)** Muscimol abolishes neuronal spiking in this injection. The top traces were recorded before muscimol injection (black) and the bottom traces were recorded 20 minutes after muscimol injection (blue). The direction tuning curve plots the mean responses of 20 trials for each direction. Direction tuning was abolished in all injections before and after training. **(D)** Average firing rate for all stimuli recorded 45 minutes after muscimol injection compared to the firing rate before injection. Firing rate went to zero in all experiments. **(E)** Average of the Local Field Potential (LFP) recorded 45 minutes after muscimol injection (blue) compared to the LFP before injection (black). The reduction of LFP amplitude is quantified as mean ± std in the rightmost panel and is similar before and after dot training (Wilcoxon rank sum test, Monkey Y: *P* = 0.81; Monkey C: *P* = 0.24).

**Figure S2, related to Figure 2.**
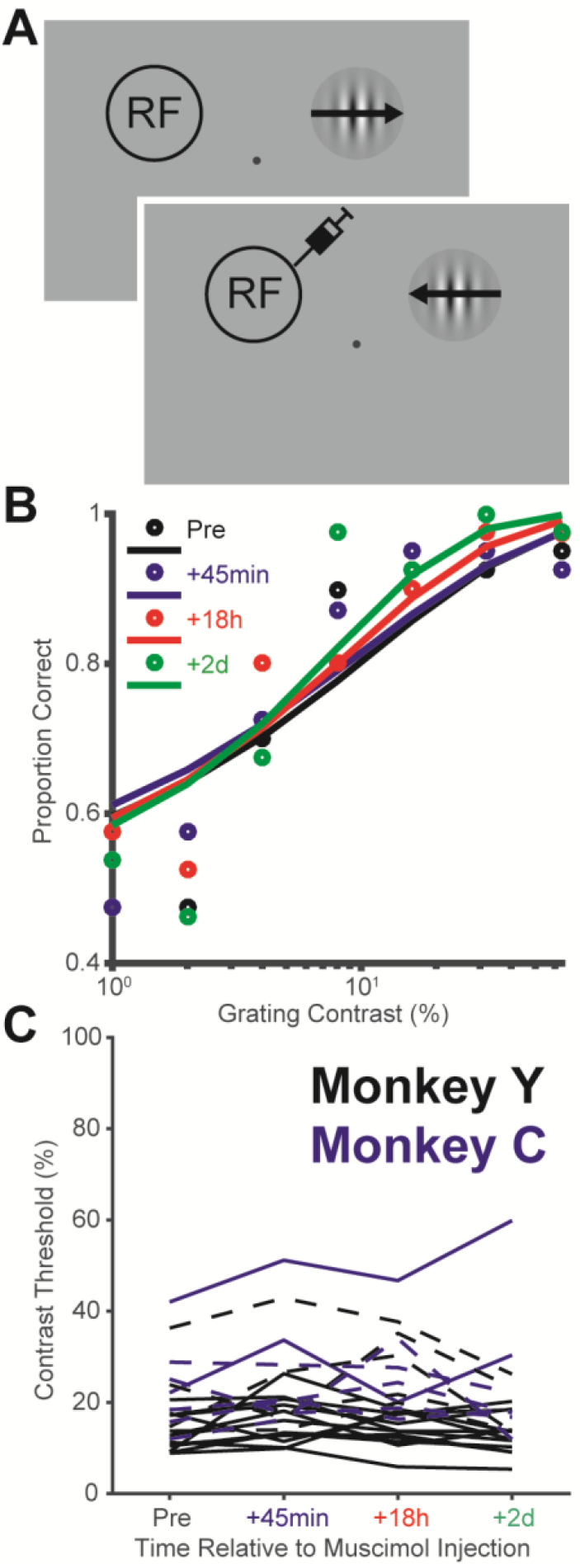
Muscimol has not effects on the control condition. **(A)** In most inactivation sessions, we included a separate control condition where the stimulus was placed in the opposite visual field. **(B)** Data from a representative experiment where the stimulus was placed in the opposite visual field. The data shows the psychophysical performance at different time points following inactivation. **(C)** Summary of inactivation effects in the control condition. The data shows the psychophysical threshold at different time points following inactivation. Solid lines are before dot training and dash lines are after dot training.

**Figure S3, related to Figure 2.**
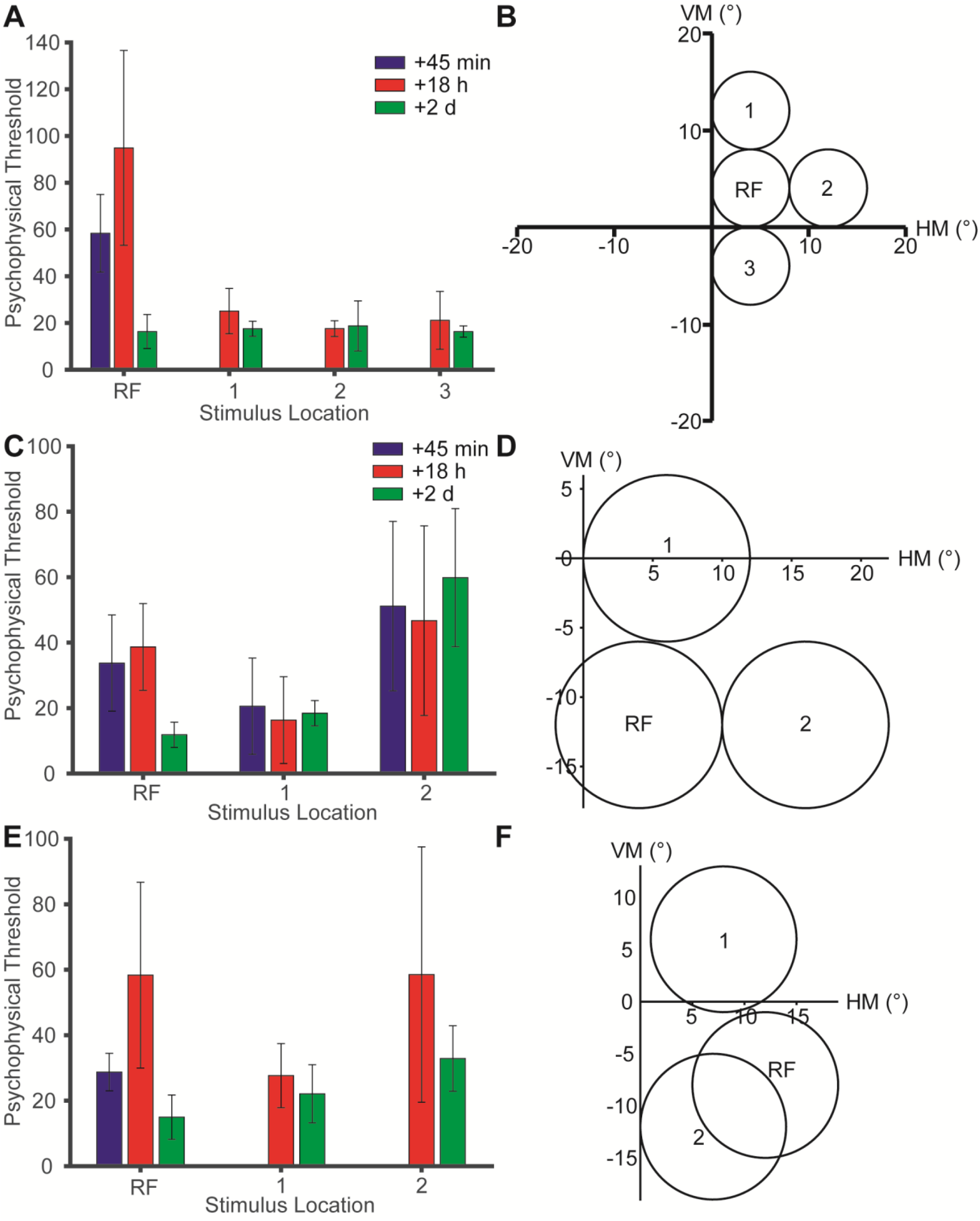
The effect of muscimol is localized in space. **(A)** Muscimol injection severely impairs motion discrimination performance for the stimulus placed in the vicinity of the region of space encoded by the MT multi-unit activity (RF). Errors bars indicate 95% confidence interval of the threshold estimates. **(B)** The dot patch was placed in the RF location of the MT neuron near the injection site, and also in the surrounding regions of visual space around the RF. The speed and size of the dot patches were tuned to the preference of the MT neuron. HM and VM indicate the horizontal and vertical meridian, respectively. **(C and D)** Another experiment showing that the psychophysical threshold is only increased at the RF location. **(E and F)** Another experiment plotted in the same format as A and B.

**Figure S4, related to Figure 2I.**
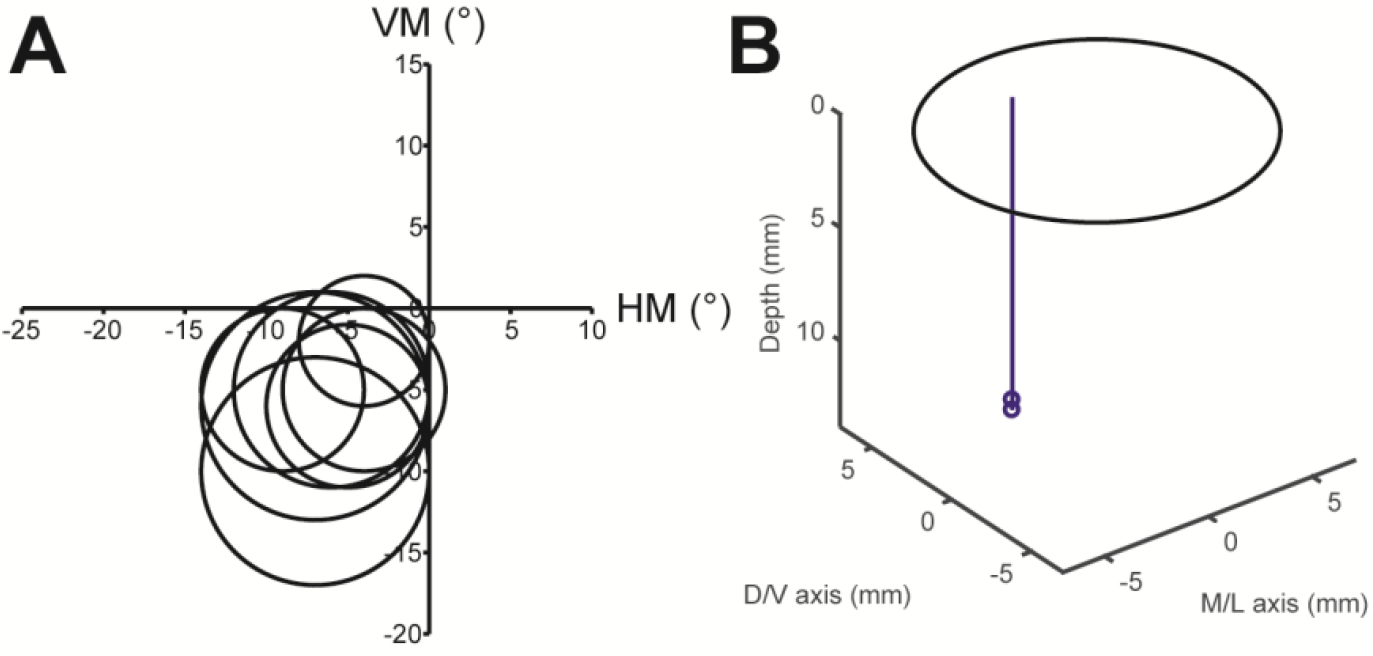
The receptive field mapping and injection sites for the experiments in Figure 2I. **(A)** Receptive field mapping for the neurons near the injection sites. HM and VM indicate the horizontal and vertical meridian, respectively. **(B)** Reconstruction of the injection sites based on grid positions and depth from the electrode microdrive. The injection sites are indicated by the circles and the electrode track is indicated by the line. The circles overlapped on 2 injection sites. M/L is medial-lateral and D/V is dorsal-ventral. The depth is zeroed at the cortical surface from a dorsal-posterior approach to MT.

**Figure S5, related to Figure 1 and 2.**
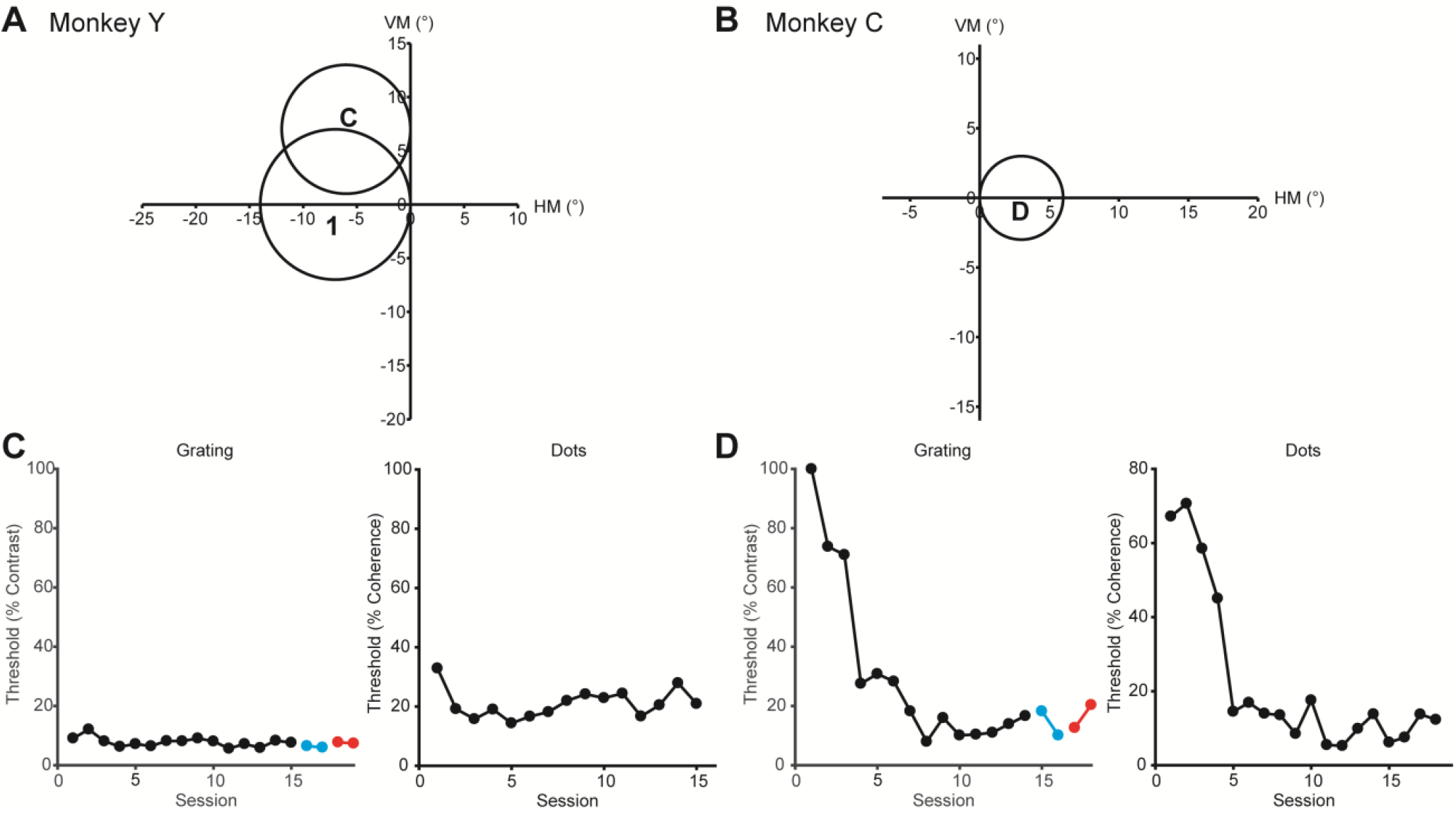
Training history of the monkeys. **(A and B)** The monkeys were trained to discriminate motion at different spatial locations. HM and VM indicate the horizontal and vertical meridian, respectively. The learning curves for Monkey Y location 1 are in Figure 1F and G. **(C)** Additional learning curves for monkey Y at the spatial location C. Once the animal generalized the task, learning at other spatial locations and stimulus parameters plateaued very quickly. The left plot is for gratings and the right plot is for dots. The blue indicates the thresholds during testing before dot training, and the red indicates the thresholds during testing after dots training. **(D)** The initial learning curves for monkey C at the spatial location D in panel B.

**Figure S6, related to Figure 2.**
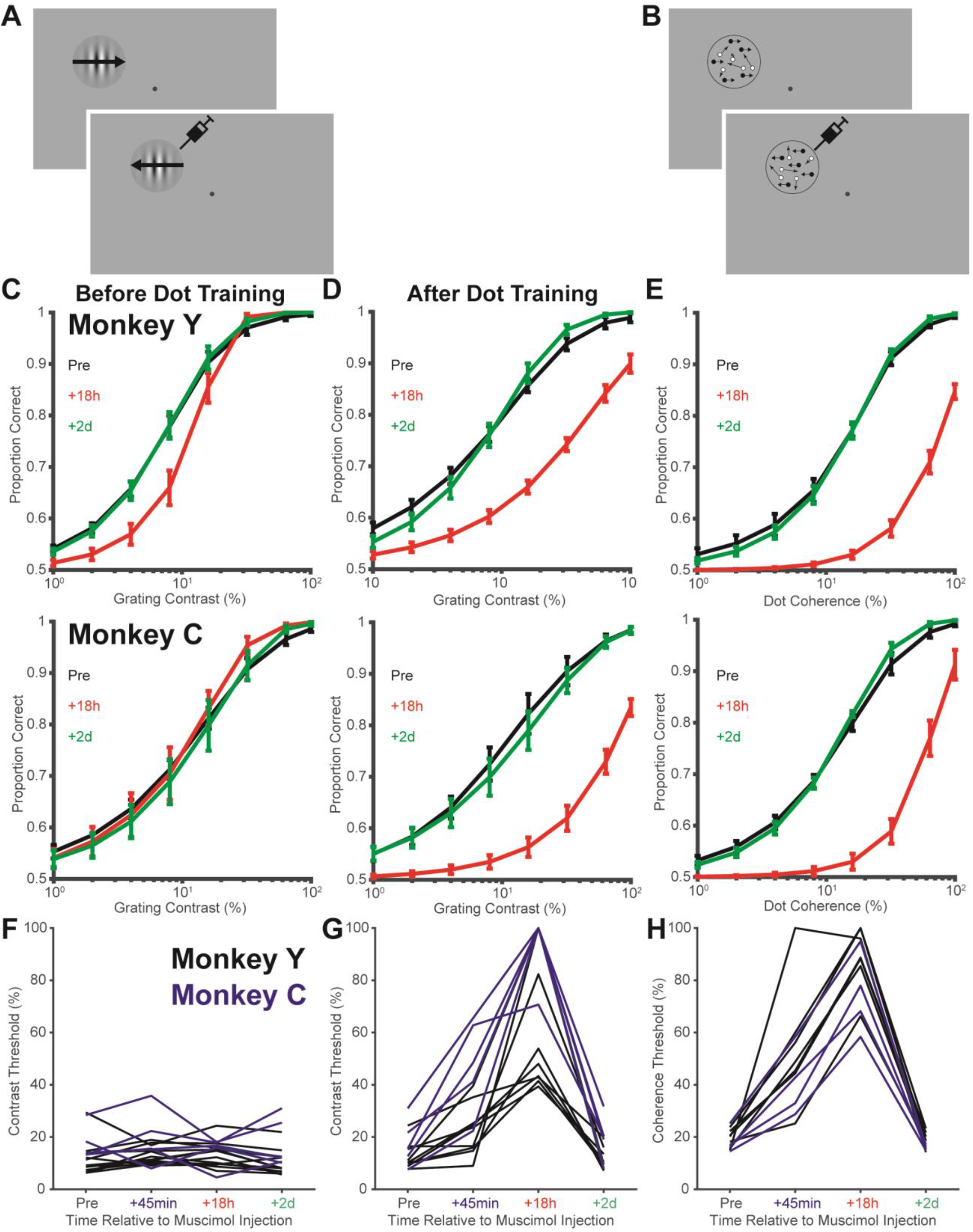
Additional psychometric functions and psychophysical testing. **(A)** Schematic illustration of the grating task. **(B)** Schematic illustration of the dot task. **(C)** Average psychometric functions before dot training measured at the pre-inactivation baseline, 18 hours after inactivation and 2 days after inactivation. **(D and E)** Average psychometric functions after dots training. **(F)** Replot of the data in Figure 2F with additional data at 45 min after inactivation. When animals were tested at 45 min after the injection, behavioral thresholds had increased by an average of only 1.9 ± 5.9% (1.9 ± 5.4% for monkey Y and 2.0 ± 7.6% for monkey C). This change was comparable to that of the variability found in the absence of inactivation, and it was not significantly different from the pre-injection baseline (Wilcoxon rank sum test; *P* = 0.19; *P* = 0.11 for monkey Y and *P* = 0.69 for monkey C). **(G and H)** Replot of the data in Figure 2G and H with additional data at 45 min after inactivation. For grating stimuli, the average threshold increase at 45 min after the injection was 17 ± 14% (10 ± 9% for monkey Y and 30 ± 13% for monkey C) (Wilcoxon rank sum test compared to the threshold change before dot training; *P* = 0.002; *P* = 0.02 for monkey Y and *P* = 0.02 for monkey C). For dot stimuli, the average threshold increase at 45 min after the injection was 31 ± 22% (35 ± 26% for monkey Y and 22 ± 9% for monkey C)

**Figure S7, related to Figure 5.**
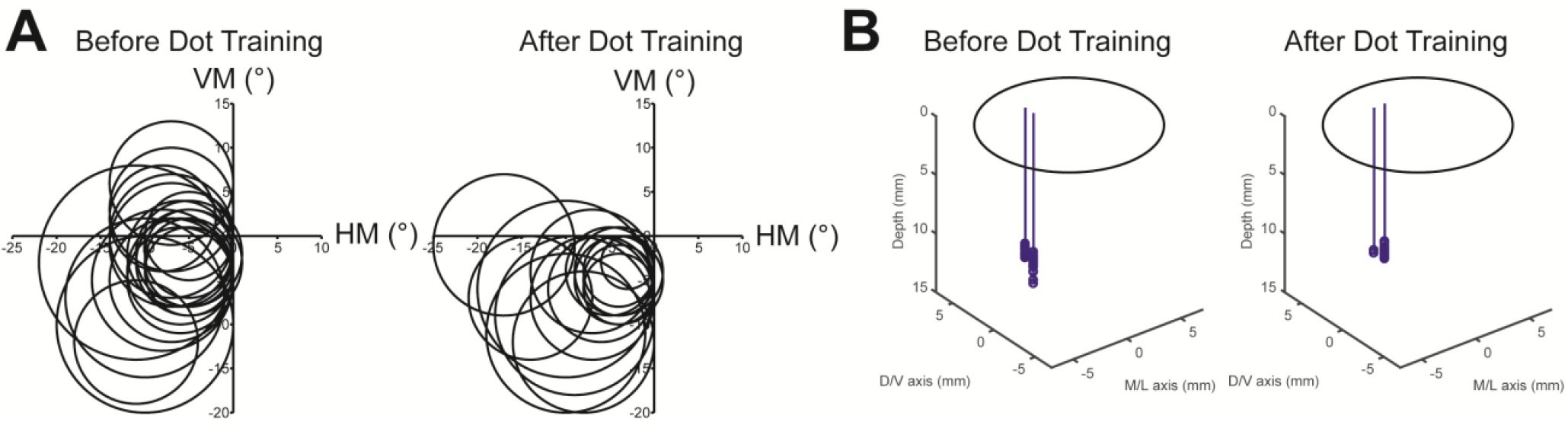
RF mapping and microstimulation sites before and after dots training. **(A)** Receptive field mapping for the neurons near the microstimulation sites. The mean RF eccentricity was 8.2 ± 3.0° before training and 9.0 ± 3.6° after training (Wilcoxon rank sum test, *P* = 0.43). The stimulus placement was based on the RF mapping. **(B)** Reconstruction of the microstimulation sites based on grid positions and depth from the electrode microdrive. The microstimulation sites are indicated by the circles and the electrode tracks are indicated by the lines. M/L is medial-lateral and D/V is dorsal-ventral. The depth is zeroed at the cortical surface from a dorsal-posterior approach to MT. The mean distance of the microstimulation sites is 13.1 ± 0.7 mm before training and 13.1 ± 0.4 mm after training (Wilcoxon rank sum test, *P* = 0.75). This is also not different from the injection sites in Figure S1B (*P* = 0.37).

